# DNA-Encircled Lipid Bilayers

**DOI:** 10.1101/285957

**Authors:** Katarina Iric, Madhumalar Subramanian, Jana Oertel, Nayan P. Agarwal, Michael Matthies, Xavier Periole, Thomas P. Sakmar, Thomas Huber, Karim Fahmy, Thorsten L. Schmidt

## Abstract

**uFig 1.**
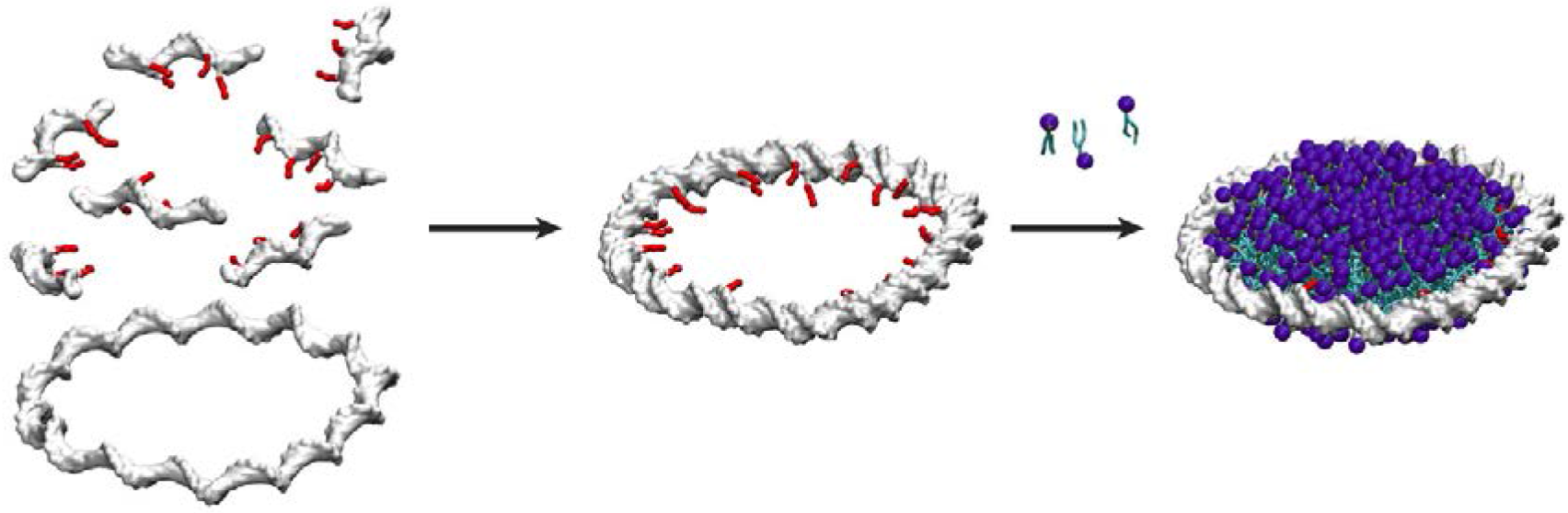

Lipid bilayers and lipid-associated proteins play crucial roles in biology. As *in vivo* studies and manipulation are inherently difficult, membrane-mimetic systems are useful for the investigation of lipidic phases, lipid-protein interactions, membrane protein function and membrane structure *in vitro.* In this work, we describe a route to leverage the programmability of DNA nanotechnology and create DNA-encircled bilayers (DEBs). DEBs are made of multiple copies of an alkylated oligonucleotide hybridized to a single-stranded minicircle, in which up to two alkyl chains per helical turn point to the inside of the toroidal DNA ring. When phospholipids are added, a bilayer is observed to self-assemble within the ring such that the alkyl chains of the oligonucleotides stabilize the hydrophobic rim of the bilayer to prevent formation of vesicles and support thermotropic lipid phase transitions. The DEBs are completely free of protein and can be synthesized from commercially available components using routine equipment. The diameter of DEBs can be varied in a predictable manner. The well-established toolbox from structural DNA nanotechnology, will ultimately enable the rational design of DEBs so that their size, shape or functionalization can be adapted to the specific needs of biophysical investigations of lipidic phases and the properties of membrane proteins embedded into DEB nanoparticle bilayers.

---

Cell compartmentalization by membranes is crucial in biology and membrane-associated proteins contribute to fundamental cellular processes in energy conversion, cell communication and signal transduction. Membrane protein function is often linked to conformational transitions which may be critically affected by lipid protein interactions.^1–5^ As *in vivo* investigations are often very challenging, such functional implications of lipid protein interactions^6^ can be more easily studied *in vitro* with artificial membrane-mimetic systems which provide a native-like lipid environment. For example, planar discoidal nanoscale lipid bilayers surrounded by amphipathic polymers^7^ or surfactant-like helical peptides^8^ have been described. Other than in spherical vesicles, these bilayers provide access to both sides of the bilayer, which may become important when studying transmembrane proteins and signal transduction. Most commonly, discoidal planar bilayers are assembled from dimeric apolipoprotein AI-derived proteins, which encircle a lipid bilayer, thereby sealing its hydrophobic rim to form nanodiscs (NDs). These membrane scaffolding proteins (MSPs) typically support lipid bilayers of 10-16 nm in diameter^9–11^. The demand for controlling the size and shape of discoidal membrane mimetics has been met by expression of MSP variants.^12^

In principle, DNA nanotechnology should allow an alternative approach to create membrane nanoparticles with defined and programmable parameters since it has proven to enable the fast *de novo* design of arbitrarily shaped structures.^13^ For large, mega-Dalton-sized structures, typically measuring tens to hundreds of nanometers, the DNA origami approach became particularly popular due to its robustness and versatility.^14–16^ For some applications, smaller structures such as tetrahedra,^17^ icosahedra,^18^ or structures from DNA minicircles (MCs)^19–23^ consisting of fewer synthetic oligonucleotides can be better suited and are more economical. Due to the full addressability of DNA structures, they can be functionalized with Å precision with a large variety of artificial elements including small molecules, fluorophores, functional groups, biomolecules or inorganic nanoparticles^24^ in a modular and programmable fashion.^16^

With the aim of leveraging these advantages of DNA nanotechnology for the design and synthesis of nanoscale discoidal lipid bilayers, we developed protein-independent DNA-encircled lipid bilayers (DEBs). In our approach, we conceptually replaced the MSP of nanodiscs by a circular double-stranded DNA minicircle (dsMC).^19–23^ To enable interactions between negatively charged hydrophilic DNA and lipids, several oligonucleotide modifications have been previously employed including cholesterol, porphyrin, phospholipid or alkyl modifications.^25^ In this way, DNA based membrane penetrating pores were created,^26,27^ or DNA Origami structures were anchored to the surface of liposomes.^28^ Furthermore, liposomes can be wrapped around DNA origami structures creating an envelope virus mimic,^29^ or liposomes can be grown inside of DNA structures,^30^ even non-spherical shapes.^31^ However, due to the tendency of lipids to form liposomes, it is challenging to stabilize a planar lipid bilayer. To date, this has not been demonstrated with a DNA scaffold.

To direct the self-assembly process towards discoidal bilayer formation, we used DNA alkylation at sites that would predictably lie on the inner circumference of the DNA minicircle. Cost-effectiveness, scalability and a linker-free chemical modification even in the middle of an oligonucleotide was achieved by alkylating phosphorothioates with alkanyl (alkyl) iodides (Figure 1 a).^34,27^ Commercially prepared oligonucleotides containing two or four internal phosphorothioates were reacted with an excess of ethyl iodide, butyl iodide or decyl iodide. As a result, the modified phosphorothioate is not charged anymore, which also increases the affinity of the modified DNA segment to the lipid bilayer rim. The respective alkylated oligonucleotides were HPLC purified and alkylation confirmed by ESI mass spectrometry (experimental details and procedures in Supporting Information).

**Figure 1.**
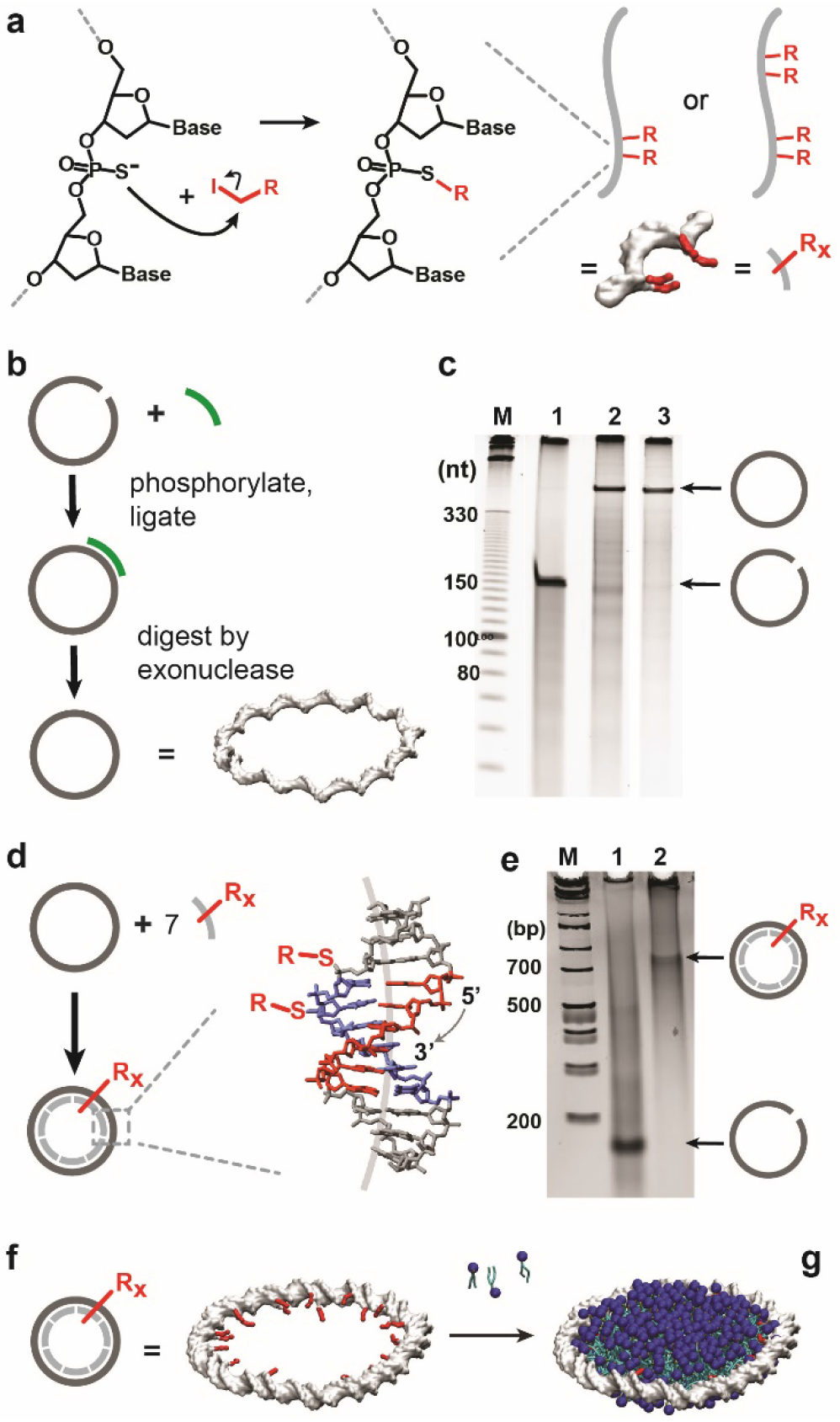
Synthesis of DEBs. a) Short oligonucleotides with two or four phosphorothioates are alkylated with alkyl-iodides (red). b) A circular single-stranded template is synthesized by enzymatic splint ligation from a long, linear oligonucleotide (grey). Residual splints (green) and linear templates are digested by exonuclease treatment. c) A denaturing PAGE gel confirms the synthesis of the single-stranded minicircle (ssMC). M, molecular size marker (nt); lane 1, linear long oligonucleotide; lane 2, ligation reaction before exonuclease treatment; lane 3, exonuclease digest. d) The ssMC is hybridized with seven alkylated oligonucleotides into double-stranded MCs (dsMC). The position of alkylations, and the intrinsic curvature of the A-tracts (grey line in the center of the helix, exaggerated) in a model of an A-tract^32,33^ (adapted from PDB structure 1FZX). Adenosines are coloured red, thymidines blue. e) Native PAGE gel. M, marker (base pairs); 1, linear long oligonucleotide; 2, the assembled dsMC complex. f) The dsMC is incubated with phospholipids (blue) to form a mature DEB (g).

In our design, the dsMC with 147 base pairs (bp) is composed of a circularized 147 nucleotide (nt) long oligonucleotide with a repetitive sequence that is hybridized to seven identical copies of the short alkylated oligonucleotide (21 nt). With a thickness of 2 nm and a rise of 0.335 nm/bp for dsDNA, their outer diameter is expected to be 16.7 nm and the inner diameter 14.7 nm. To define the inside and outside of the desired toroidal dsMC, the sequence was designed with 14 intrinsically curved A-tracts,^32,35^ as depicted in Figure 1 d.^33^ First, a single-stranded minicircle (ssMC) was prepared from one long oligonucleotide (147 nt) by enzymatic splint ligation (Figure 1 b). Residual linear long oligonucleotides, splints and linear side products were enzymatically removed by a treatment with exonuclease I/III (Figure 1 b-c). Next, the ssMCs were hybridized with the alkylated oligonucleotides by slow cooling. Excess oligonucleotides were removed by ultrafiltration and the double-stranded minicircles (dsMCs) analyzed by native agarose gel electrophoresis (Figure 1 e), atomic force microscopy (AFM) and transmission scanning electron microscopy (tSEM). Finally, the alkylated dsMCs were filled with a lipid bilayer (Figure 1 f-g). Similar to nanodiscs formation with MSPs, removal of detergent from a detergent-solubilized mix of the alkylated dsMCs and stoichiometric amounts of phospholipids (MC:lipid was 1:450) led to the self-assembly of the components into nanoscale discoidal particles (Figure 2).

**Figure 2.**
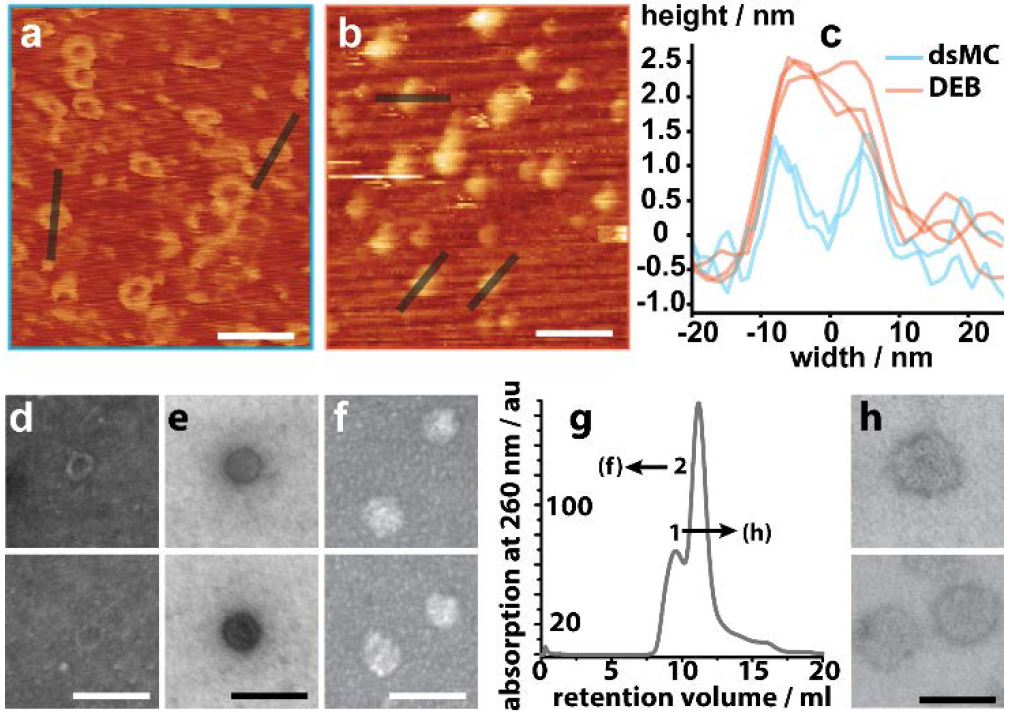
Analysis of dsMCs and DEBs. a) an AFM image of empty dsMCs (R = 4 butyl) and corresponding DEBs (b). c) height profiles. d) tSEM images of empty dsMCs. e) tSEM image of a DEB with 14 ethyl modifications (positively stained). f) tSEM image of a DEB with 28 decyl modifications. g) elution profile of a size exclusion chromatography (SEC) run of a DEB preparation. Fraction 1 contains dimeric DEBs (h), fraction 2 monomers (f). Scale bars, 50 nm.

AFM imaging of the dsMCs and DEBs revealed a doubling of the height of the empty DNA minicircles from ~1.3 nm to ~2.5 nm due to the addition of the lipid bilayer (Figure 2 a-c). Absolute diameters or heights of soft, compressible biomolecules can usually not be measured with standard AFM imaging due to the mechanical deformation during scanning caused by the AFM tips and additional deformations caused by interactions with highly charged surfaces. As a result, both the DNA (actual thickness = 2 nm) and the DMPC bilayer (thickness = 4-5 nm) appear thinner than in force-free environments. The tSEM images (Figure 2 d, f) also confirm the presence of a lipid bilayer in the DEBs. Short (14 ethyl) and longer alkyl chains (28 decyl), produced DEBs (Figure 2 e,f), but DEBs with longer alkyl chains formed with higher yields (Figure 3 a-c).

**Figure 3.**
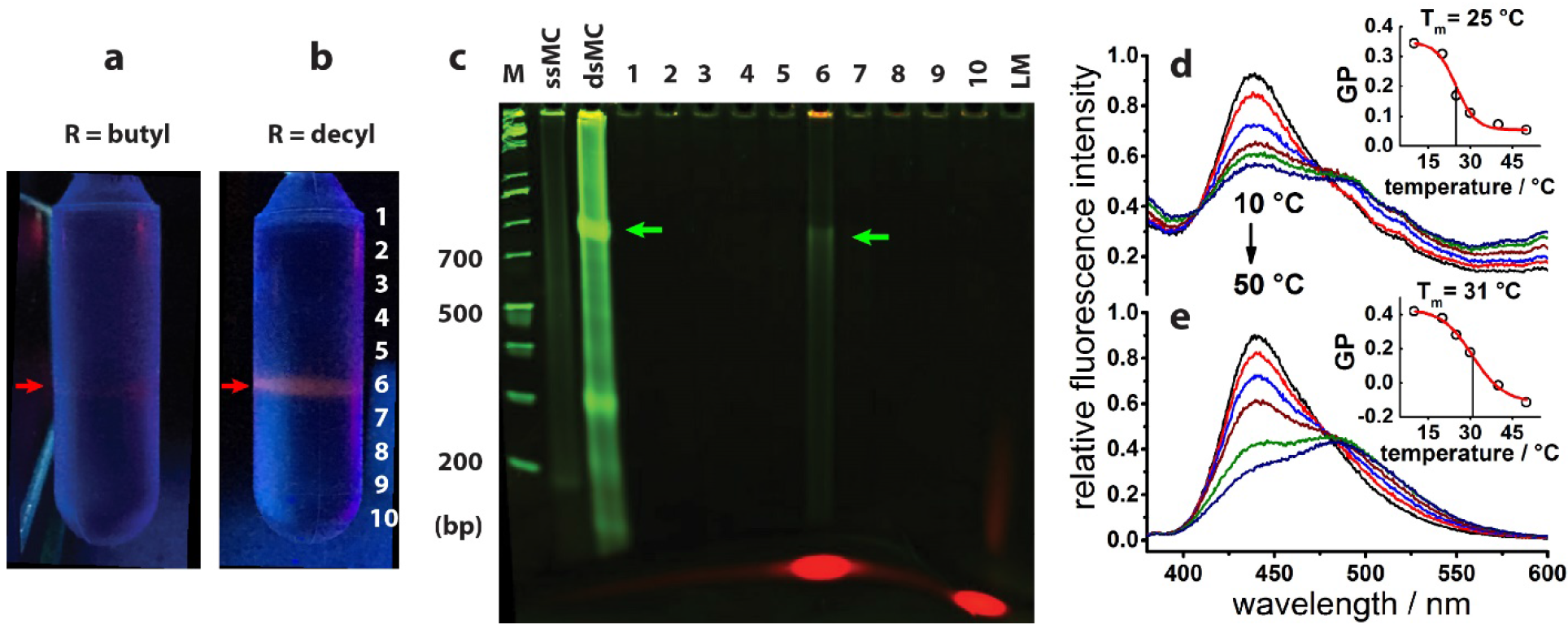
Analysis of DEBs. Gradient ultracentrifugation results of Rhodamine-PE-containing DEBs with 28 butyl groups (a), and 28 decyl groups (b) that were analyzed by native SDS PAGE (c). LM = control lipid mix. d-e) Lipid phase transition of DEBs. Emission spectra of LAURDAN (λ_exc_ = 340 nm) in DMPC-filled DEBs carrying two ethyl groups per hybridized 21-mer (d) and in MSP-based NDs (e) recorded at 10 °C, 20 °C, 25 °C, 30 °C, 40 °C and 50 °C. Inserts show the temperature dependence of the generalized polarization, GP = (I_440_ - I_490_) / (I_440_ + I_490_) which reveals the gel to liquid phase transition^41^ (T_m_: transition midpoint temperature).

DEBs were purified either by size exclusion chromatography (Figure 2 g) or by ultracentrifugation, where DEBs formed one sharp band that contained both lipids and DNA. In syntheses of ssMCs at high concentrations, dimeric ligation products occurred as a side product (Figure S1). They were separated by SEC from the monomers and exhibited twice the circumference of monomers (Figure 2 g,h). This result demonstrates that the DEB approach can extended to enable designs with custom sizes.

Next, we compared the thermotropic phase transition of DMPC in conventional MSP-based NDs with that of DEBs by using the emission of the lipophilic dye LAURDAN as a sensor of lipid order.^36^ The dye binds at the sub-headgroup region of lipids. Its excited state energy depends on the dipolar relaxation processes in its environment leading to a red shift of LAURDAN fluorescence with increasing hydration that accompanies the gel to liquid phase transition of the lipid. Therefore, the phase transition can be monitored by the relative intensity difference measured at two LAURDAN emission wavelengths, i.e. the so-called general polarization (GP).^36^ The midpoint temperature T_m_ for the gel to liquid transition of DMPC in DEBs was 25 °C and agrees with literature data on DMPC vesicles,^37^ whereas DMPC in NDs showed a slightly higher T_m_ (31 °C) as reported.^38^ However, the change of the GP value in DEBs was only ~50% of that in NDs transition, which may indicate restricted lipid mobility at the alkylated DNA-lipid interface. Upon doping of the DMPC bilayer with the cationic lipid DMTAP, no phase transition was observed (Figure S2), which we attribute to the additional electrostatic interactions of the positively charged head groups with the dsMC, and to a preferential binding of LAURDAN at the DNA lipid interface (Supporting Information).

We next prepared a coarse grain molecular dynamics (CGMD) simulation of a DEB using the MARTINI force field (Figure 4).^39,40^ In agreement with experimental results, a DEB was stable over a 5 microsecond duration with the 4 nm thick DMPC bilayer encircled by the 2 nm thick dsDNA rim. The interaction of the alkyl chains (Figure 4a-b, red) with the lipid bilayer could also be clearly observed, demonstrating the effectiveness of our modification strategy.

**Figure. 4.**
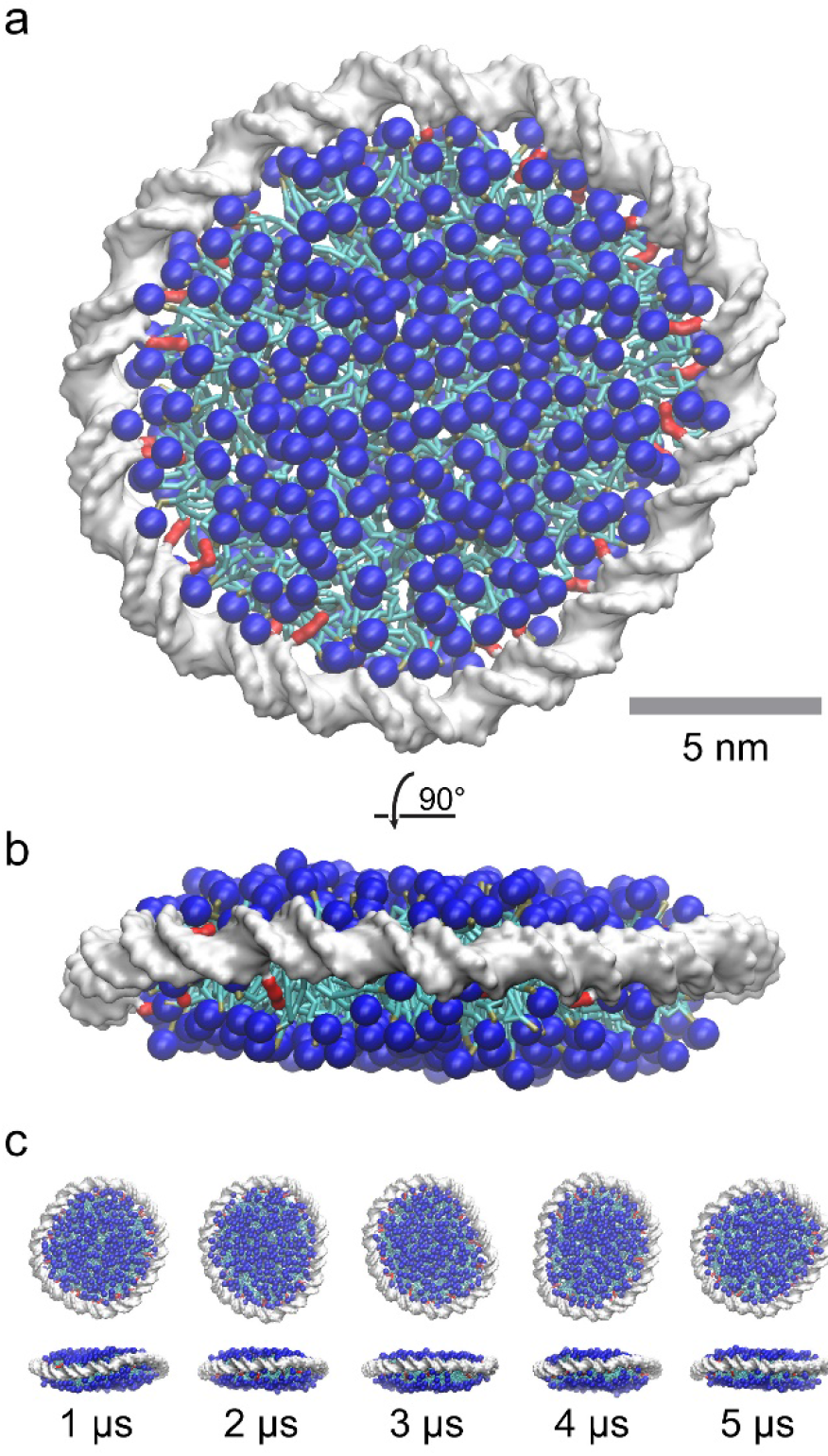
Coarse grain molecular dynamics model of a DEB composed of a 147 bp dsMC with 28 dodecyl groups and 434 DMPC lipids. DNA is white, alkyl chains red, DMPC head groups blue. a) View down the membrane normal of a structure at the end of the 5 microseconds long trajectory. b) rotated 90 degrees. c) Snapshots after 1-microsecond intervals.

In summary, we report the formation of DEBs as a novel strategy to prepare nanoscale discoidal bilayer structures encapsulated by an alkylated dsMC. The difficulty in preparing nanoscale membrane mimetics is to prevent formation of vesicles, which are the preferred state of lipid bilayers. Thus far, only the shaping of spherical vesicles with DNA structures has been reported. With the DEB technology we have realized for the first time the use of DNA to assemble planar lipid bilayers with dimensions that are comparable to protein-stabilized NDs. In contrast to the DNA origami approach, which requires hundreds of oligonucleotides and expensive single-stranded scaffold strands, the minimalistic DEB design can be accomplished with only two commercial synthetic oligonucleotides and facilitates upscaling. Only one oligonucleotide has to be chemically modified, and the chosen alkylation of phosphorothioate is among the most economical and scalable modification approaches. We have demonstrated a high alkylation density of up to two DNA backbone alkylations per helical turn without additional linkers, to stabilize the lipid bilayer. In addition to the demonstrated ease of DEB size variation, the core structure of DEBs is provided by a covalently circularized ssMC. The proposed DEB design is thus inherently independent of more sophisticated biochemistry required for improving stability and mono-dispersity of protein-based scaffolds via circularization.^41^ We anticipate that further developments of the DEB technology will provide a nanoscale membrane mimetic that profits from the attractive links to DNA technology towards higher order assemblies and spatial arrangements of lipid bilayers for studying their interactions with membrane-associated proteins.

## ASSOCIATED CONTENT

### Supporting Information

The following files are available free of charge. Supporting information (PDF) containing materials and methods, and Supporting Figures.

### Funding Sources

This work was funded by the DFG through a staring grant of cfaed to T.L.S, a fellowship of the DIGS-BB to K.I. and a networking grant from the Helmholtz-Association to K.F. and M.S. Support was also provided to T.H. from the Robertson Therapeutic Development Fund.

### Notes

K.F, KI and T.L.S. have filed a provisional patent application.

## ACKNOWLEDGMENT

We thank Prof. Zhang and Dr. Reddavide (B CUBE) for help with HPLC-ESI MS. We are grateful to Lisa Nucke for many helpful discussions and to Jenny Philipp for technical support. We acknowledge the use of the imaging facilities in the Dresden Center for Nanoanalysis (DCN), the skillful advice from Dr. Löffler and the use of the AFM of Prof. Mertig (TU Dresden).

