## Supplementary Materials for "DNA-Encircled Lipid Bilayers" (from another version of this preprint: version 1, 2018-03-21)

Enzymes and respective buffers were purchased from New England Biolabs. Oligonucleotides were synthesized by Integrated DNA Technologies and chemicals were purchased from Sigma-Aldrich unless stated otherwise. Lipids in chloroform solution were purchased from Avanti Polar Lipids. Denaturing polyacrylamide gel electrophoresis (PAGE) gels (15% TBE urea gels), SYBR Gold gel stain and Pierce detergent removal columns were purchased from Thermo Fisher Scientific. Ladders for the PAGE were either 10 bp ladder (Invitrogen) or O'GeneRuler$^{TM}$ 1kb plus ladder (Thermo Fisher).

##### Oligonucleotide sequences

short oligonucleotide with 2 phosphorothioate (PTO) groups: 5'-TTT TTC ACA CTT T*TT* CAC ACT-3'
short oligonucleotide with 4 PTO groups: 5'-TT*T T*TC ACA CTT T*TT* CAC ACT-3'
(asterisks indicate the locations of backbone phosphorothioates)
long oligonucleotide: 5'-(TGT GAA AAA AGT GTG AAA AAG)$_7$-3'
splint: 5'-CTT TTT TCA CAC TTT TTC AC-3'

### Methods

All the experiments were performed at room temperature unless stated otherwise.

##### Alkylation of short oligonucleotides

The chemical modification of the phosphorothioated oligonucleotides was conducted by following a modified protocol of Gut and Beck [1]. Phosphorothioate–modified oligonucleotides (8.3 nmol, 21 nt) were reacted with 245 nmol alkyl-iodide in 90% DMF (Alfa Aesar) and 10% 30 mM Tris-HCl pH 8.0 (1250 µL). The mixture was incubated at 65 °C for 3 h. The excess of the organic solvent was removed by gel permeation using NAP-25 columns (GE healthcare). The collected fraction was purified by reversed phase HPLC using a Dionex (ICS-5000+ TC) system with a MultoKrom 100-5 C18 column (flow rate 1 mL/min) using the following gradient starting points (difference to 100%: triethyl ammonium acetate (TEAA; 20 mM; pH 8)):

0 min, 3% acetonitrile (ACN)

10 min, 10 % ACN

25 min, 21% ACN

30 min, 100% ACN

35 min, 100% ACN

40 min, 3% ACN

The retention time for the starting compound (phosphorothioated oligonucleotide) was 8.1 min, whereas the retention time for the alkylated oligonucleotides was 15.2 min (2 ethyl groups per short oligonucleotide), 19.0 min (4 butyl groups per short oligonucleotide) and 26.1 min (4 decyl groups per short oligonucleotide). The last of typically 3-5 closely spaced peaks contained the intended product with all alkyl chains attached; the earlier peaks contained not fully alkylated products. In some samples, two closely spaced peaks with the same mass were observed. We attribute this to different possible stereoisomers of the phosphorothioates. In this case, both peaks were combined. Reaction yields were typically 30-40 % as determined by the integrals of the HPLC traces at 260 nm.

**Mass spectrometry**

The mass of the alkylated oligonucleotides was determined from the collected HPLC fractions with a combined HPLC-MS system (Waters ACQUITY UPLC S4 with a Ultra Performance/Waters ACQUITY TQ Detector). The masses of the functionalized oligonucleotides were determined by deconvolution of the peaks with multiple charges.

R = 2 ethyl, calculated 6362 Da, measured: 6361

R = 2 butyl expected: 6418 Da; measured: 6416

R = 4 butyl, calculated 6562 Da, measured 6559

**ss DNA MC design and preparation**

The long oligonucleotide (147 nt, IDT Ultramer) and the splint (19 nt) were designed with the OligoAnalyzer Tool (idtdna.com) disregarding potential sequences with strong secondary structures. The long oligonucleotide (3.33 µM, 100 pmol) was hybridized to the splint (66.67 µM, 2 nmol) in 1X ligase buffer (total volume: 30 µL) using a 12 min thermal annealing program (80 °C, 1 min, 80°C – 25 °C -5 °C/min, 25 °C hold).

The annealed mixture was diluted twofold in 1X ligase buffer and treated with 1 µL polynucleotide kinase and 1 µL T4 ligase for 1 h at room temperature, after which another 0.5 µL of each enzyme were added and the reaction mixture was incubated for one more hour at 30 °C. The mixture was purified with Zymo Oligo Clean & Concentrator columns (Zymo research) according to the manufacturer's protocol and the DNA was recovered in 40 µL of water. The non-circular byproducts or linear oligonucleotides were digested in 1X exonuclease I buffer supplemented with 1X exonuclease III buffer, and 1.5 µL of exonuclease I and 1.5 µL exonuclease III (total volume: 55 µL) for 3 h at 37 °C. The removal of the enzymes and buffer exchange were performed with Zymo Oligo Clean & Concentrator. ssMCs were recovered in 20 µL of water and analyzed by denaturing PAGE. The yield of ssMCs was 80%.

**Denaturing PAGE**

For the denaturing PAGE analysis, 15% gels were cast with 1X TBE buffer containing 6 M urea and run in 1x TBE buffer. 9 µL of sample solution were mixed with 9 µL of 2X gel loading dye (7 M urea, 89 mM Tris, 89 mM boric acid, 2 mM EDTA, pH 8.0, 12% ficoll, 0.01% bromophenol blue, 0.02% xylene cyanole). As a reference, 2 µL of 10 bp DNA ladder (Thermo Scientific) were added.

The gels were run at 200 V, ~55 °C for 45 min inside an isolating Styrofoam box filled with ~2 L of hot tap water, after which they were stained and imaged the same way as described native PAGE (see below).

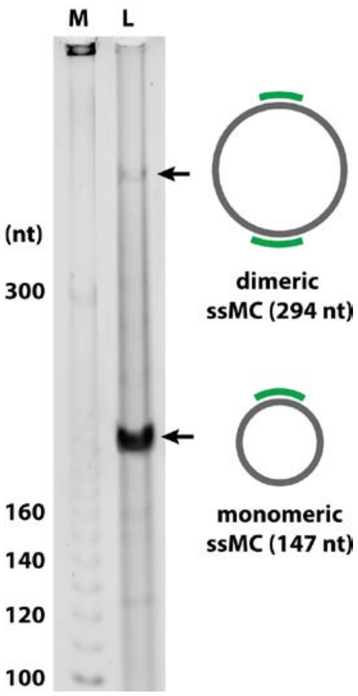


***Figure S1:*** *Denaturing PAGE gel (6 %) of a splint ligation experiment for the synthesis of ssMC after exonuclease treatment. For entropic reasons, circularizations are favored over polymerizations. In some experiments, dimeric single-stranded minicircles were observed as side products. These dimeric ssMC can hybridize with the alkylated strands to produce dimeric dsMCs and lead to DEBs with twice the circumference (see Figure 2 h). Note that the electrophoretic mobility of the ssMC is different from the one in Fig. 2c. In general, circulated oligonucleotides have a lower mobility than their linear equivalents, but the extent of this difference is a function of the concentration of the PAGE gel. In Fig. 2c, a 15% gel was used, whereas a 6% gel is shown that was chosen to increase the resolution for the high molecular weight ssMC dimer. The splints (green) are only added to the scheme for clarity; they are digested during the exonuclease step.*

**dsDNA MC formation**

The ds DNA MCs were formed by hybridization of the ssDNA ring (7 µM) with complementary alkylated oligonucleotides (70 µM) using a 12 min thermal annealing program (80 °C, 1 min, 80°C – 25 °C -5 °C/min, 25 °C hold) in 12 mM $MgCl_2$ and 5 mM TE buffer (50 µL). The excess of the complementary oligonucleotides was removed by ultrafiltration as described below. The purified ds MCs were analyzed by native PAGE.

**Purification of ds MCs with ultrafiltration columns**

Passivation of the ultrafiltration filter (10 kDa MWCO, Amicon Ultra, AMD Millipore) membranes was carried out by incubating them in 400 µl of 1X buffer H (50 mM HEPES pH 7.4, 200 mM $Na_2SO_4$ and 5 mM $MgSO_4$) for 30 min, after which the buffer was passed through the filters at 14,000 rfc for 2 min. The assembled ds DNA MC solution was added to the passivated filters and washed two times with 400 µL of 1X buffer H at 14,000 rfc for 13 min. After the final wash, the filter was reversed, placed in a fresh tube, and centrifuged at 1,000 rfc for 2 min. The collected purified dsMC solution was used for further characterization and reactions.

**Native PAGE**

For the native PAGE analysis, 10% gels were cast with 1X TBE buffer containing 12 mM $MgCl_2$. Running buffer contained 1X TBE buffer and 12 mM $MgCl_2$. 15 µL of sample solution were mixed with 3 µL of 6X gel loading dye (50% glycerol, 5 mM Tris-HCl, 1 mM EDTA, 12 mM $MgCl_2$, 0.25% bromophenol blue and 0.25% of xylene cyanol). Electrophoresis was performed at 65 V for 12 h at room temperature. As a reference, 2 µL of 1 kb plus DNA ladder (Thermo Scientific) were added. After the electrophoresis, the gel was removed from the gel cassette and stained for 10 min with 1X SYBR Gold in 50 mL 1X TBE supplemented with 5% ethanol and imaged with a Typhoon FLA 9500 gel scanner (GE Healthcare) using the excitation wavelength of 488 nm suitable for SYBR Gold stained gels.

**DEB formation and purification**

14:0 TAP (1,2-dimyristoyl-3-trimethylammonium-propane), 14:0 Rhodamine-PE (1,2-dimyristoyl-*sn*-glycero-3-phosphoethanolamine-N-(lissamine rhodamine B sulfonyl)) and DMPC (1,2-dimyristoyl-sn-glycero-3-phosphocholine were mixed in a 9.9:1.4:88.7 molar ratio to reach 112.3 µL total volume. The chloroform was evaporated using an argon line for 2 h, followed by drying under vacuum for the next 2 h. The lipid film was dissolved in 90 µl of 60 mM sodium cholate in buffer H (final concentration: 1.5 mM) and sonicated five times for 1 min. Next, 30 µL of the ds DNA MCs were added (2.4 µM) to obtain a ratio of 450 lipids per dsMC and the mixture was incubated for 1 h at room temperature. The detergent was removed with Pierce detergent removal columns according to manufacturer's protocol and eluted in 120 µL buffer H. DEBs were further purified in 50mM HEPES (containing 100 mM $Na_2SO_4$ and 8 mM $MgSO_4$) on a Superdex 200 Increase column (GE Healthcare) at a flow rate of 0.75 ml/min using an Akta Avant system (GE Healthcare).

**Ultracentrifugation**

Unpurified DEB solution (120 µL) was mixed with 630 µL of 45% iodixanol in 1X buffer H and placed at the bottom of a centrifuge tube (6.0 mL, 13 x 64 mm, Beckman Coulter). Six additional layers of iodixanol solution (26%, 22%, 18%, 14%, 10%, 6%, and 2%) in 1X buffer H (750 µL) were added to the centrifuge tube. The tube was spun in a SW-41-Ti rotor (Beckman Coulter) at 41,000 rpm = 288,000 rfc and 4 °C for 6 h, after which the contents were fractionated (500 µL per fraction) and analyzed by $MgCl_2$-supplemented SDS PAGE (see below).

**$MgCl_2$-supplemented SDS PAGE**

$MgCl_2$-supplemented SDS PAGE gels (10%) were cast with 0.13 M Tris-HCl buffer pH 8.8, 12 mM $MgCl_2$ and 0.05% SDS. 6.25 mM Tris-HCl buffer with 12 mM $MgCl_2$ and 0.05% SDS was used as a running buffer. 10 µL of sample solution (DEBs carrying 2 ethyl chains per 21-mer) were mixed with 10 µL of SDS gel loading dye (0.2 M Tris HCl, 0.1% SDS, 20% Glycerol and 0.25% of bromophenol blue). Electrophoresis was performed at 65 V for 12 h. As a reference, 3 µL of 1 kb plus DNA ladder (Thermo Scientific) were added. The gel was imaged with a Typhoon FLA 9500 gel scanner using the excitation wavelength of 575 nm suitable for 14:0 Rhodamine PE signal. After the first imaging, the gel was post-stained with SYBR Gold and imaged the same way as native PAGE.

**tSEM characterization**

Carbon-coated TEM grids (400 mesh copper, carbon on Formvar, Science Services Munich) were plasma-treated for 15 s. 4 µL of the sample solution were applied on the TEM grid and incubated for 5 min. The excess solution was removed from the grid with a filter paper wick.

Next, 5 µL of a 1% uranyl formate solution was applied for 90 s to stain the DEBs, and the solution was removed with a filter paper. The samples were scanned on a Gemini SEM500 (Zeiss) SEM/STEM system operated at 10 kV.

**AFM imaging**

70 µL of a 0.01% poly-L-ornithine solution was applied on a freshly cleaved mica plate. After 1 min the plate was washed with water and blown dry using compressed air. A small circle (diameter ca. 3 mm) was drawn using a permanent marker. 3 µl of sample were applied into the center of the circle and incubated for 1 min. Next, the mica plate was covered with 70 µl of imaging buffer (120 mM $NiCl_2$, 5 mM Tris*HCl pH 8.0). The scans are performed using a Cypher ES AFM (Asylum research) using BL-AC40TS tips (BioLever Mini, Olympus). The AFM was operated in AC mode, at a scanning frequency of 0.5 Hz and a set point of 300 mV. Height profiles were obtained with Gwyddion software.

**Expression and purification of MSP1D1**

The standard expression of MSP1D1[2] was adapted from literature. [3] Briefly, the MSP1D1-pET28a plasmids were grown overnight in BL21Gold (DE3) cells (Agilent Technologies) at 37 °C in double strength YT medium containing 50 µg/mL kanamycin. After induction with 0.3 mM isopropyl-β-D-thiogalactopyranoside, the temperature was decreased to 28 °C. The cells were harvested 4 h later, frozen in liquid nitrogen, and stored at -80°C until further use. For purification, cells were resuspended in buffer A (50 mM Tris–HCl, 200 mM NaCl, pH 7.4) containing protease inhibitors (Roche Applied Science) and lysed twice by sonicating at 30% amplitude for two minutes with repeated 30 s pulse and 30 s pause. Cell debris was removed by centrifugation (15,000 g, 70 min, 4°C). Imidazole was added to the supernatant to a final concentration of 25 mM. The sample was loaded onto a Ni-NTA column (GE-Healthcare), equilibrated with buffer B (buffer A containing 25 mM imidazole). The column was washed with buffer B. Finally, the protein was eluted by gradient elution using increasing concentrations of imidazole (280 mM, 500 mM and 1 M; one CV each). Fractions containing MSP1D1 were identified by SDS-PAGE. A desalting step with PD10 columns (GE Healthcare) equilibrated with buffer A was performed to remove imidazole. The desalted MSP1D1 sample was concentrated using Vivaspin4 columns (Sartorius) and the final protein concentration measured by absorbance at 280 nm using a calculated extinction coefficient of

21,430 $M^{-1}$ $cm^{-1}$ and a molecular weight of 24,793 kDa (ProtParam, ExPASy). Purified MSP1D1 was frozen in liquid nitrogen and stored at -80 °C until further use.

**Assembly of MSP-based lipid nanodiscs**

The protocol of phospholipid nanodisc assembly was adapted from the original publication. [4] A completely dried DMPC (1,2-dimyristoyl-*sn*-glycero-3-phosphocholine, Avanti Lipids) lipid film was solubilized in buffer A, containing cholate as detergent twice the concentration of the lipid, and sonicated until a clear solution was obtained. The respective lipid/sodium cholate solution and MSP1D1 were mixed to yield a final concentration of 12 mM lipid and 0.2 mM MSP1D1 (DMPC:MSP1D1 = ratio 60:1). For fluorescence measurements, LAURDAN was added in a ration 1:200 to DMPC, respectively. The mixture was incubated for 1 h at 25 °C. The detergent was removed with Detergent Removal spin columns (Pierce). The approximate size and the homogeneity of the DMPC-filled nanodiscs were verified by size exclusion chromatography.

**Fluorescence measurements of LAURDAN-containing DEBs and vesicles**

Fluorescent measurements were performed a Fluorolog 3 FL3-11 (HORIBA Europe GmbH, Oberursel). Samples containing 1-(6-(dimethylamino)-2-naphthalyl)-1-dodecanon (LAURDAN) in 100 µL cuvettes were excited at 340 nm (10 nm bandwidth) and the general polarization (GP) calculated[5] from emissions at 440 and 490 nm as GP = (I440-I490) / (I440 + I490) if not stated otherwise. Scan speed was 100 nm $min^{-1}$ and 900 V photomultiplier voltage was chosen. The emission spectrum was averaged from 4 scans. A thermostatted cuvette holder connected to a circulating water bath was used to record temperature-dependent spectra after three minutes of thermal equilibration. Figure S2 shows the temperature-dependent LAURDAN emission spectra of DEBs with twenty-eight ethyl or butyl chains stabilizing a bilayer composed of a 1:10 DMTAP:DMPC mixture. In both cases, the main emission band was blue-shifted and broadened (with lobes at 410 and 433 nm) and additional peaks occurred at 495 and 519 nm. Whereas a blue shift of LAURDAN emission upon association with cationic lipids has been reported[6], the additional spectral features indicate the partitioning of the dye into different local lipid environments of the DEBs. Remarkably, the spectral deviations from the normal LAURDAN emission are specific to the presence of the DNA scaffold and are absent in vesicles of the identical lipid-mixture (Fig. S2 a, insert) where the temperature-dependent GP values follow a typical phase transition (Fig.

S2 b). Therefore, we tentatively assign the additional spectral features to LAURDAN populations that occupy regions at the DNA lipid interface where hydrophobic mismatch between the cationic lipid and the alkylated DNA may have been compensated by the dye, for example, between the alkylation sites. At such sites, the LAURDAN hydration may not be linked to a thermotropic lipid transition which nevertheless may have occurred in the center of the DEB. Irrespective of the detailed physical origin of the emission shift, the data demonstrate that vesicle formation was negligible during detergent removal for DEB assembly, since the temperature dependency of GP, which is typical of non-DNA-bounded DMTAP/DMPC vesicles, was negligible in DEBs.

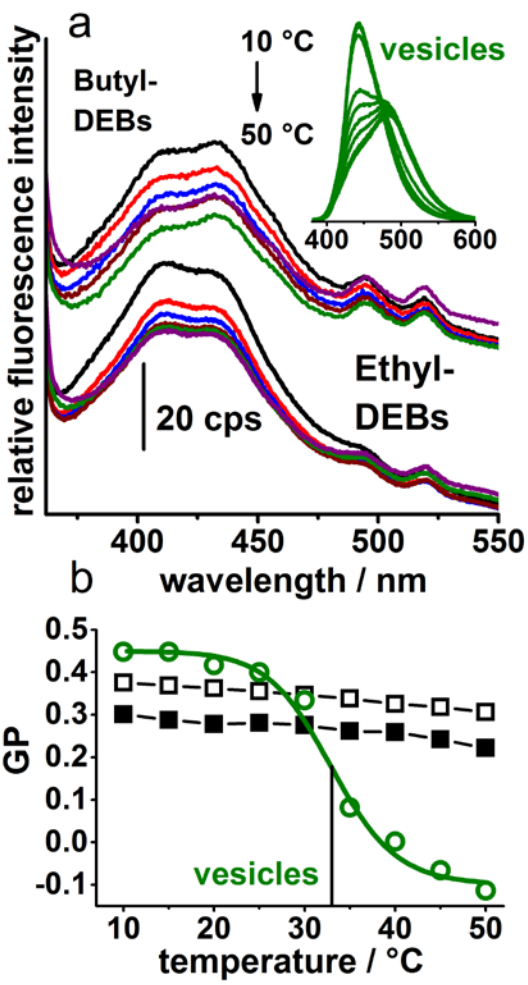


***Figure S2: Phase transition of DMTAP-doped DMPC in DEBs and in vesicles.*** *a) Emission spectra of LAURDAN ($\lambda_{exc}$ = 340 nm) in DMTAP:DMPC (1:10) recorded at 10 °C, 20 °C, 25 °C, 30 °C, 40 °C and 50 °C with DEBs carrying four ethyl or butyl chains per 21-mer (cps: counts per second). Insert: LAURDAN emission from the identical lipid mixture in vesicles. b) The generalized polarization (GP) calculated from the intensities at 435 nm and 495 nm indicates a marginal phase transition in both ethylated (open squares) and butylated (filled squares) DEBs. The GP value determined for vesicles comprised of the same DMTAP/DMPC mixture (circles) evidences a phase transition with $T_m$= 33 °C.*

#### Coarse-grain molecular dynamics calculations

**Models.** A circular DNA model in a standard internal coordinate representation (with tilt, roll, twist, shift, slide, and rise inter base pair step parameters) was generated with a modified version of the vdna-plugin-2.2[7] using vmd-1.9.3.[8] We modified the plugin to include the repeating 21-mer sequence motif. We converted the internal coordinates into an atomistic model using the *rebuild* module of the x3dna-2.3[9] package with non-standard residues encoding the chain breaks between the 21mers and the alkylated phosphorothioates. For the coarse-grained models, we built a separate model with unmodified bases to facilitate the conversion to the Martini model, which then was modified to include chain breaks and alkylations.

This atomistic model of the ring/DNA complex was converted into a MARTINI-v2.2 coarse grained model using the martinize script. [10] The MARTINI force field for lipids[11] and its extension to DNA[12] were used in combination with the ElNeDyn approach[13] to maintain the structure of the DNA backbone. A tight elastic network was applied as suggested by Uusitalo *et al*. [12] This CGMD approach is well suited to study complex biological membranes. [14]

Inspired by a recent study by Maingi *et al.,*[15] we used conventional MARTINI beads and parameters to describe the S-alkyl chains modifications to the DNA rings. The S atom was mapped to the phosphate group to which it was chemically linked in the atomistic model. This new backbone bead was assigned a P5 type and the charge was set to zero. The beads of the alkyl chain were modeled by a C2 bead. All additional bonds were given a 4.5 nm reference value and a 5000 $kJ/mol·nm^{-2}$. Bond angles of the first bead of the chain relative to the backbone beads (previous and following bead, at 105 and 145 degrees, respectively, and with a force constant of 50 $kJ/mol·rad^{-2}$) were used to restrict the orientation of the chain. The remaining bond angles were set to 180 degrees with a 25 $kJ/mol·rad^{-2}$ force constant. Improper dihedrals were used to force planarity of the backbone bead anchor of the alkyl chain, its neighbors in the chain and the first bead of the chain. This precaution prevents back flips of the chain.

**Simulations.** All molecular dynamics simulations were performed using the GROMACS simulation package version 5.1. [16] Conventional simulation setups associated with the MARTINI[10, 17] were used. These include a 20 fs time step for production run, a 0.9 nm cutoff and 500 $kJ/mol·nm^{-2}$ force constant for ElNeDyn, and non-bonded interactions (van der Waals and electrostatic) cutoff at 1.1 nm and shifted to zero using the potential-modifier implementation in GROMACS. A relative dielectric screening constant of 15 and the Verlet neighbor search[18] were used. The DNA, the membrane bilayer (DMPC), and the aqueous phase (water plus ions when present) were coupled independently to an external temperature bath at 300 K using Berendsen[19] and Bussi[20] thermostats ($\tau_T$=0.5 ps) for equilibration and for production, respectively. The pressure was weakly coupled[19] to an external bath at 1 bar using a relaxation time of 2 ps following a semi-isotropic pressure scheme. Each system was run for 5 microseconds.
